## Supplemental information for "Acquisition of auditory discrimination mediated by different processes through two distinct circuits linked to the lateral striatum"

**Figure supplements legends**

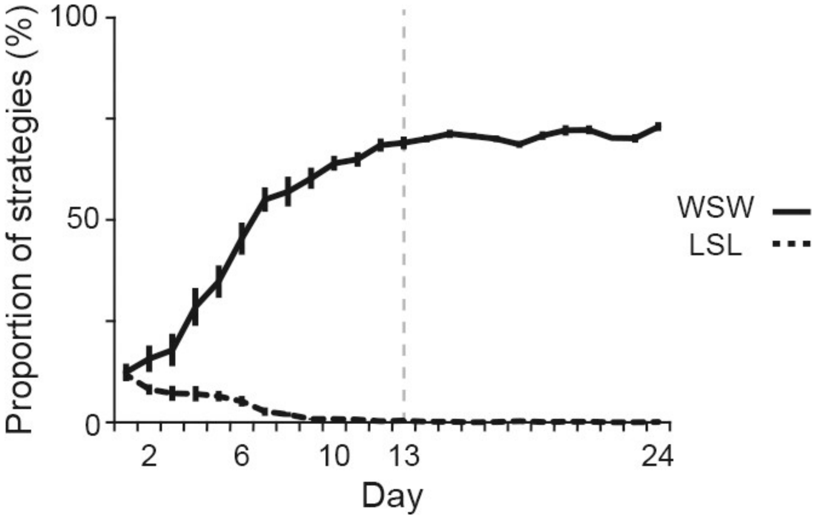


**Figure 1–figure supplement 1**. **Proportions of behavioral strategies during auditory discrimination learning.** Proportions of the win-stay-win and lose-shift-lose strategies in intact rats ($n$ = 14) shown in Figure 1. Vertical dashed line indicates the day of transition between the acquisition and learned phases. Data are indicated as the mean ± s.e.m. Abbreviations: WSW, win-shift-win; and LSL, lose-shift-lose.

**Figure 2–table supplement 1. Task-related brain activity during the acquisition of auditory discrimination**.

| Learning days | Brain regions | Laterality | T-value (peak) | Volume* ($mm^3$) |
|---|---|---|---|---|
| | *Cerebral cortex* | | | |
| | Entorhinal cortex | R | 5.52 | 4.75 |
| | | L | 8.48 | 11.18 |
| | Perirhinal cortex | L | 4.71 | 0.20 |
| | Primary somatosensory cortex, jaw region | R | -3.93 | 0.84 |
| | | L | -4.89 | 0.29 |
| | Primary somatosensory cortex, barrel field | R | -5.36 | 0.84 |
| | | L | -4.97 | 0.20 |
| 2 | *Midbrain* | | | |
| | Inferior colliculus | L | -4.88 | 0.47 |
| | *Medulla* | | | |
| | Inferior olive | R | 6.89 | 1.03 |
| | *Cerebellum* | | | |
| | Copula of the pyramis | L | 5.01 | 0.59 |
| | Crus 2 of the ansiform lobule | L | 6.51 | 0.85 |
| | Cerebellar lobule 2 | R | -4.83 | 0.57 |
| | Cerebellar lobule 7/8 | R | -5.57 | 0.37 |
| | *Cerebral cortex* | | | |
| | Primary motor cortex | L | 6.48 | 3.59 |
| | Secondary motor cortex | R | 5.80 | 1.32 |
| | | L | 5.77 | 3.59 |
| | Perirhinal cortex | L | 4.75 | 0.25 |
| | *Basal ganglia* | | | |
| | Anterior dorsolateral striatum | R | 6.70 | 1.99 |
| | | L | 4.76 | 0.63 |
| | Posterior ventrolateral striatum | R | 4.51 | 0.21 |
| | Nucleus accumbens | R | 4.50 | 0.44 |
| | Substantia nigra pars compacta/reticulata | L | 5.61 | 1.54 |
| 6 | *Hippocampus* | | | |
| | Hippocampus CA1 field | L | 6.26 | 1.02 |
| | *Thalamus* | | | |
| | Ventral posterolateral thalamic nucleus | L | 5.14 | 0.83 |
| | Medial geniculate body | L | 5.23 | 0.49 |
| | *Medulla* | | | |
| | Ventral cochlear nucleus | R | -7.16 | 16.45 |
| | *Cerebellum* | | | |
| | Crus 1 of the ansiform lobule | R | -4.79 | 16.45 |
| | | L | -5.86 | 0.64 |
| | Paraflocculus | R | -7.38 | 16.45 |
| | | L | -5.49 | 4.87 |
| | *Cerebral cortex* | | | |
| | Agranular insular cortex posterior part | R | 4.25 | 0.15 |
| | | L | 4.60 | 0.73 |
| | Entorhinal cortex | R | 8.05 | 4.12 |
| | Orbitofrontal cortex | R | -7.24 | 1.81 |
| 10 | *Basal ganglia* | | | |
| | Posterior ventrolateral striatum | R | 7.30 | 0.43 |
| | | L | 5.57 | 2.57 |
| | Substantia nigra pars compacta/reticulata | R | 4.35 | 1.40 |
| | *Amygdala* | | | |
| | Central amygdaloid nucleus | R | 6.91 | 0.66 |

| | | | | |
|---|---|---|---|---|
| | *Thalamus* | | | |
| | Medial geniculate body | R | 5.41 | 0.24 |
| | | L | 4.39 | 0.11 |
| | Mediodorsal thalamic nucleus | L & R | -6.82 | 5.59 |
| | Ventral anterior thalamic nucleus | L & R | -6.08 | 5.59 |
| | Habenular nucleus | L | -5.02 | 1.00 |
| | *Midbrain* | | | |
| | Dorsal cochlear nucleus | L & R | 8.47 | 13.12 |
| | *Medulla* | | | |
| | Pontine reticular nucleus | L | -5.47 | 1.28 |
| | Dorsal tegmental nucleus | L | -5.92 | 13.41 |
| | *Cerebellum* | | | |
| | Crus 1 of the ansiform lobule | L | -4.94 | 0.28 |
| | Cerebellar lobule 3/4/6 | L | -7.88 | 13.41 |
| | Cerebellar lobule 10 | L | -5.33 | 13.41 |
| | Paraflocculus | R | -9.41 | 6.65 |
| | | L | -6.20 | 9.41 |
| | *Cerebral cortex* | | | |
| | Prelimbic cortex | L & R | -6.26 | 2.19 |
| | Agranular insular cortex | R | 6.24 | 22.36 |
| | | L | 7.10 | 25.85 |
| | Cingulate cortex/Secondary motor cortex | L & R | -7.40 | 117.86 |
| | Primary somatosensory cortex, barrel field | R | -4.59 | 0.22 |
| | | L | -5.40 | 117.86 |
| | Primary somatosensory cortex, trunk region | R | -4.22 | 0.24 |
| | Retrosplenial cortex | L & R | -8.19 | 117.86 |
| | Temporal association cortex | R | 4.10 | 2.58 |
| | | L | 6.44 | 25.85 |
| | Entorhinal cortex | L | -5.89 | 0.44 |
| | Claustrum | R | 5.74 | 0.96 |
| | *Basal ganglia* | | | |
| | Dorsomedial striatum | L | -6.94 | 117.86 |
| | *Hippocampus* | | | |
| | Hippocampus CA1 field | R | -4.46 | 0.24 |
| | | L | -8.59 | 117.86 |
| | *Thalamus* | | | |
| | Mediodorsal thalamic nucleus | L & R | -8.98 | 117.86 |
| 24 | Laterodorsal thalamic nucleus | L & R | -7.54 | 117.86 |
| | *Midbrain* | | | |
| | Dorsal cochlear nucleus | L & R | 7.73 | 37.60 |
| | *Pons* | | | |
| | Pontine nuclei | L & R | -6.21 | 1.34 |
| | Locus coeruleus | R | -4.97 | 0.31 |
| | *Medulla* | | | |
| | Medial mammillary nucleus | L & R | 8.47 | 37.60 |
| | *Cerebellum* | | | |
| | Crus 1 of the ansiform lobule | L | 5.78 | 3.90 |
| | Crus 2 of the ansiform lobule | R | 6.06 | 1.94 |
| | | L | 5.41 | 3.90 |
| | Cerebellar lobule 3/4/6 | R | -7.82 | 4.69 |
| | Cerebellar lobule 10 | R | -4.91 | 0.26 |
| | Paramedian lobule | R | 4.36 | 0.23 |
| | Paraflocculus | R | -7.48 | 117.86 |
| | | L | -9.90 | 9.17 |
| | Lateral cerebellar nucleus | R | -7.74 | 117.86 |
| | | L | -6.46 | 9.17 |
| | Medial vestibular nucleus | L | -5.37 | 0.49 |

Rats ($n$ = 14) were trained using the single lever press task, followed by the auditory discrimination task. $^{18}$F-FDG-PET scanning was conducted on Day 4 of the single lever press task, and then during a series of auditory discrimination task on Days 2, 6, 10, and 24. The regional brain activity related to behavioral tasks was calculated with a voxel-based statistical parametric analysis that compared $^{18}$F-FDG images on the single lever press task with those on either day (Days 2, 6, 10, and 24) of the discrimination task. Task-related brain regions with greater (T-value > 0) or smaller (T-value < 0) activity are summarized. The brain activity was increased or decreased in various brain regions during the progress of auditory discrimination. Especially, the increased activity was observed in the medial geniculate body, which sends ascending information to the auditory cortex and also receives feedback corticothalamic projections (***Lee et al., 2013***). However, there was no significant changes in the brain activity in the auditory cortex, being consistent with the results of a previous work reporting that the auditory cortex does not show the correlation to the pure-tone discrimination task (***Gimenez et al., 2015***). R and L indicate right and left hemispheres, respectively. The value of 3.8 was used as the threshold corresponding to the $p < 0.001$ (uncorrected) threshold. *Original volume.

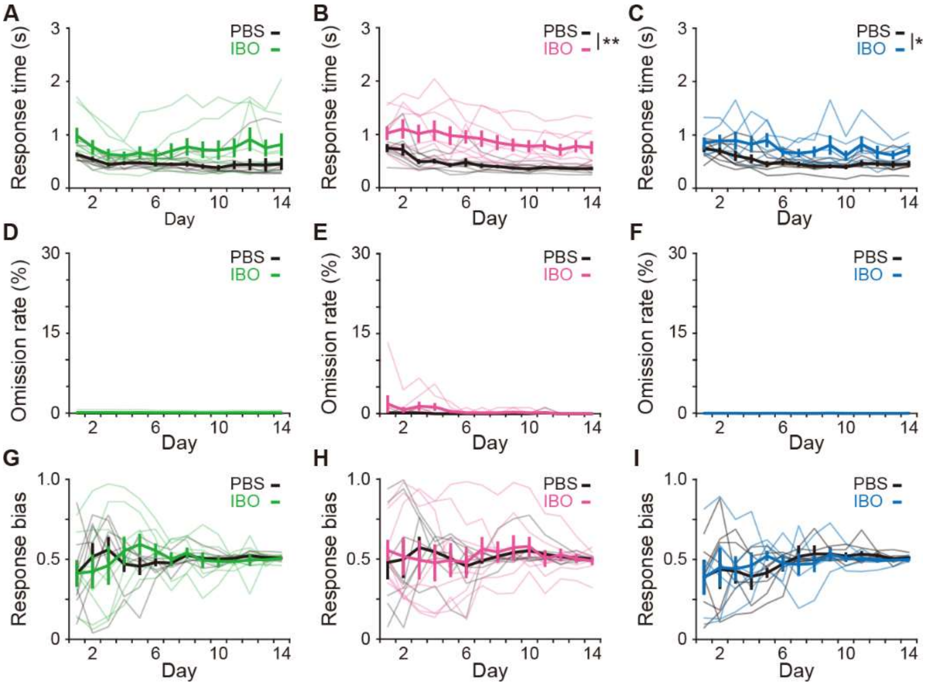


**Figure 3–figure supplement 1. Effects of excitotoxic lesion on the reaction time, omission ratio, and response bias.** (A-C) Response time (A, two-way repeated ANOVA, day, $F[2.728,38.193] = 2.867$, $p = 0.054$, group, $F[1,14] = 3.562$, $p = 0.080$, day × group, $F[2.728,38.193] = 1.137$, $p = 0.343$ for the aDLS; B, day, $F[2.890,40.456] = 6.230$, $p = 0.002$, group, $F[1,14] = 13.566$, $p = 0.002$, day × group, $F[2.890,40.456] = 0.685$, $p = 0.561$ for the pVLS; C, day, $F[3.123,34.358] = 4.659$, $p = 0.007$, group, $F[1,11] = 6.474$, $p = 0.027$, day × group, $F[3.123,34.358] = 1.127$, $p = 0.353$ for the DMS). (D-F) Omission rate (D, two-way repeated ANOVA, day, $F[1.066,14.927] = 1.038$, $p = 0.330$; group, $F[1,14] = 1.041$, $p = 0.325$, day × group, $F[1.066,14.927] = 0.800$, $p = 0.393$ for the aDLS; E, day, $F[1.384,19.372] = 1.483$, $p = 0.247$, group, $F[1,14] = 2.297$, $p = 0.152$, day × group, $F[1.384,19.372] = 1.140$, $p = 0.320$ for the pVLS; F, day, $F[3.119,34.306] = 2.398$, $p = 0.083$, group, $F[1,11] = 0.950$, $p = 0.351$, day × group, $F[3.119,34.306] = 0.951$, $p = 0.429$ for the DMS). (G-I) Response bias (G, two-way repeated ANOVA, day, $F[2.573,36.027] = 0.748$, $p = 0.512$, group, $F[1,14] = 0.008$, $p = 0.929$, day × group, $F[2.573,36.027] = 0.853$, $p = 0.459$ for the aDLS; H, day, $F[2.523,35.317] = 0.430$, $p = 0.699$, group, $F[1,14] = 0.008$, $p = 0.931$, day × group, $F[2.523,35.317] = 0.349$, $p = 0.756$ for the pVLS; I, day, $F[2.429,26.718] = 1.286$, $p = 0.297$, group, $F[1,11] = 2.4 \times 10^{-5}$, $p = 0.996$, day × group, $F[2.429,26.718] = 0.348$, $p = 0.749$ for the

DMS). Data are indicated as the mean ± s.e.m., and individual data are overlaid. *$p < 0.05$ and **$p < 0.01$.

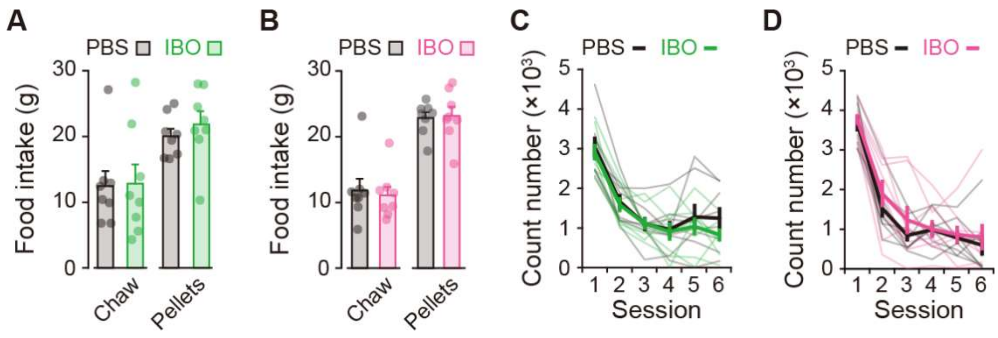


**Figure 3–figure supplement 2. Effects of excitotoxic lesion on feeding behavior and locomotion.** (A and B) Amounts of food intake (laboratory chow or reinforcement pellets) for 60 min (A, unpaired Student's t-test, laboratory chow, $t[14] = 0.101$, $p = 0.921$; reinforcement pellets, $t[14] = 0.796$, $p = 0.439$ for the aDLS; B, laboratory chow, $t[14] = 0.345$, $p = 0.735$; reinforcement pellets, $t[14] = 0.201$, $p = 0.844$ for the pVLS). (C and D) Time course of the number of beam breaks for a 10-min block in the open field (C, two-way repeated ANOVA, session, $F[2.214,30.994] = 34.082$, $p = 6.8 \times 10^{-9}$, group, $F[1,14] = 0.530$, $p = 0.479$, session × group, $F[2.214,30.994] = 0.348$, $p = 0.730$ for the aDLS; D, session, $F[5,70] = 79.020$, $p = 2.1 \times 10^{-27}$, group, $F[1,14] = 0.559$, $p = 0.467$,session × group, $F[5,70] = 0.382$, $p = 0.860$ for the pVLS). Data are indicated as the mean ± s.e.m., and individual data are overlaid.

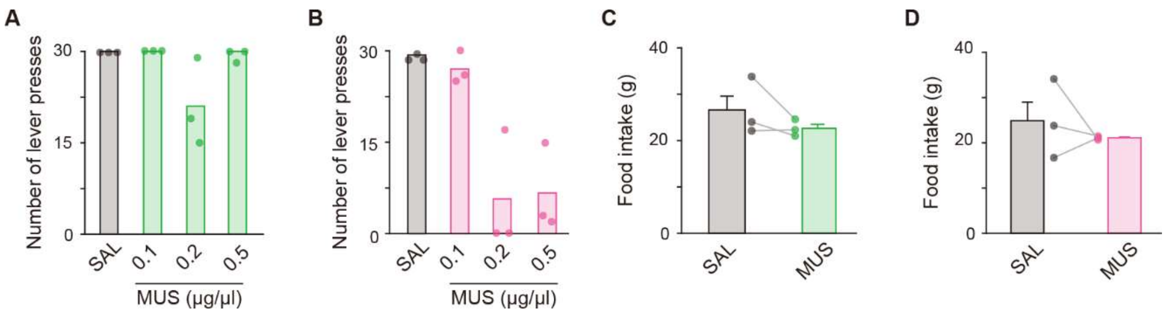


**Figure 4–figure supplement 1. Effects of bilateral MUS injections into the aDLS and pVLS on the single lever press task and feeding behavior.** Rats (*n* = 3 for each group) received a bilateral injection of solution containing MUS or SAL into the aDLS and pVLS, and then conducted the behavioral task. (A and B) Number of single lever presses in group injected with SAL or different doses of MUS into the aDLS (A, one-way repeated ANOVA, $F[3,6] = 4.315$, $p = 0.061$) or the pVLS (B, one-way repeated ANOVA, $F[3,6] = 27.236$, $p = 6.8 \times 10^{-4}$). (C and D) Amount of food intake for 60 min after the SAL and MUS (0.1μg/μL) injections into the aDLS (C, paired Student's t-test, $t[2] = 1.450$, $p = 0.284$) or the pVLS (D, paired Student's t-test, $t[2] = 0.726$, $p = 0.543$). Data are indicated as the mean ± s.e.m., and individual data are overlaid.

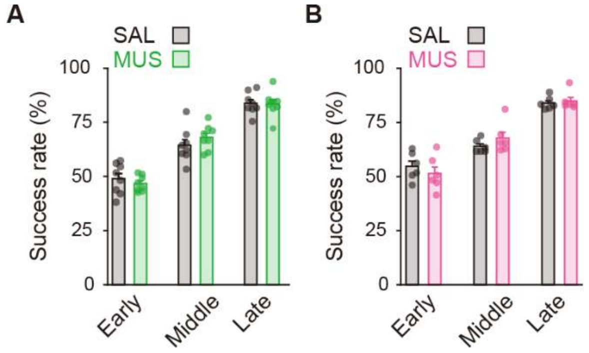


**Figure 4–figure supplement 2. Performance of the auditory discrimination task on Day N-1 of rats used for the injection into striatal subregions.** The success rate in the groups used for the aDLS injection (A, unpaired Student's t-test, early, $t[10.437] = 0.800$, $p = 0.441$; middle, $t[14] = 1.064$, $p = 0.305$; late, $t[14] = 0.118$, $p = 0.907$) or the pVLS injection (B, unpaired Student's t-test: early, $t[10] = 0.789$, $p = 0.448$; middle, $t[10] = 1.224$, $p = 0.249$; late, $t[10] = 0.489$, $p = 0.635$). Data are indicated as the mean ± s.e.m., and individual data are overlaid.

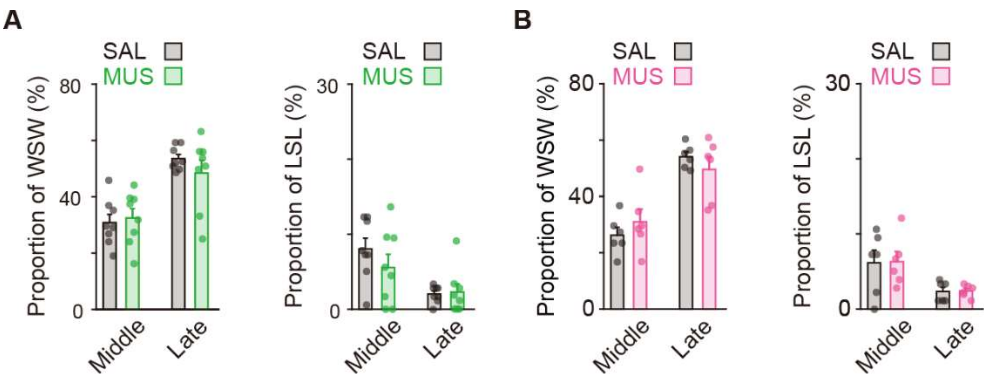


**Figure 4–figure supplement 3. Proportion of the behavioral strategy on Day N-1 of rats used for the striatal injection.** The strategy proportion in the groups used for the aDLS injection (A, unpaired Student's t-test, win-shift-win: middle, $t[14] = 0.376$, $p = 0.713$; and late, $t[8.479] = 1.078$, $p = 0.311$. lose-shift-lose: middle, $t[14] = 1.084$, $p = 0.297$; and late, $t[14] = 0.229$, $p = 0.822$) or the pVLS injection (B, unpaired Student's t-test, win-shift-win: middle, $t[10] = 0.912$, $p = 0.383$; and late, $t[6.408] = 0.942$, $p = 0.380$. lose-shift-lose: middle, $t[10] = 0.075$, $p = 0.942$; and late, $t[8.267] = 0.129$, $p = 0.900$). Data are indicated as the mean ± s.e.m., and individual data are overlaid. Abbreviations: WSW, win-shift-win; and LSL, lose-shift-lose.

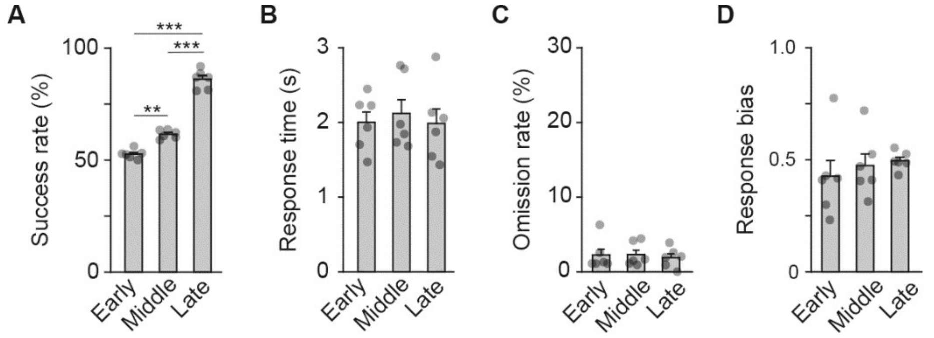


**Figure 5–figure supplement 1. Behavioral performance of rats used for the multi-unit recording experiment.** Behavioral parameters at the early, middle, and late stages. Success rates (A, one-way repeated ANOVA, $F[2,10] = 213.186$, $p = 6.3 \times 10^{-9}$, *post hoc* Bonferroni test; early vs. middle, $p = 0.003$; early vs. late, $p = 5.7 \times 10^{-5}$; middle vs. late, $p = 4.9 \times 10^{-5}$). Response time (B, one-way repeated ANOVA, $F[2,10] = 0.379$, $p = 0.694$). Omission rate (C, one-way repeated ANOVA, $F[2,10] = 0.163$, $p = 0.852$). Response bias (D, one-way repeated ANOVA, $F[2,10] = 0.600$, $p = 0.567$). Data are indicated as the mean ± s.e.m., and individual data are overlaid. $^{**}p < 0.01$ and $^{***}p < 0.001$.

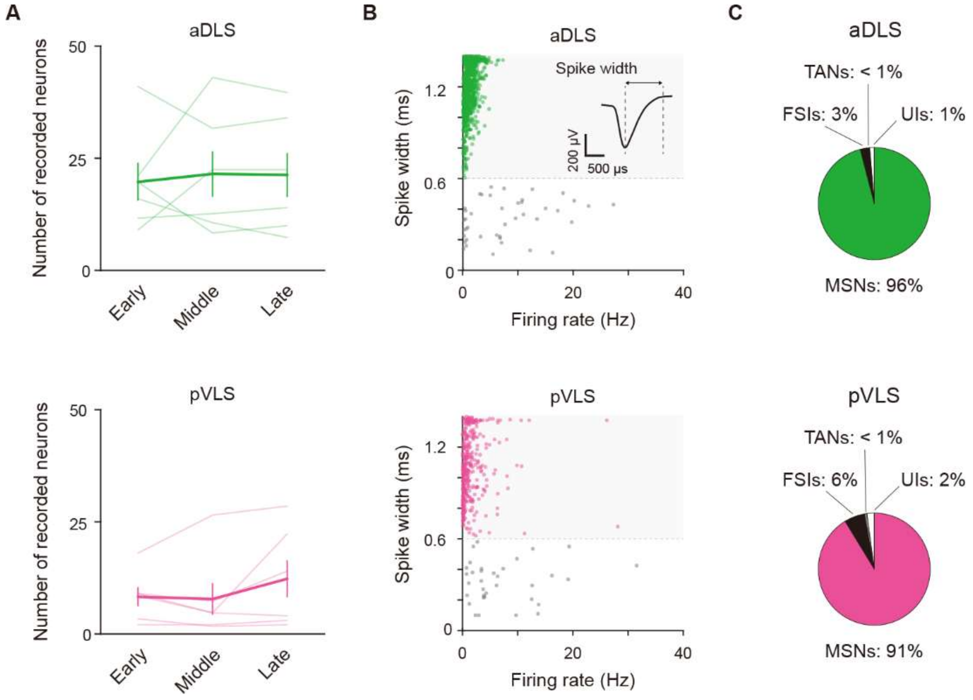


**Figure 5–figure supplement 2. Properties of identified neurons in the multi-unit recording experiment.** (A) Averaged number of well-isolated neurons for every stage in individual rats (one-way repeated ANOVA, aDLS, $F[1.045,5.226] = 0.098$, $p = 0.777$; pVLS, $F[2,10] = 1.855$, $p = 0.207$). (B) Scatter plots of spike width and firing rate of aDLS and pVLS neurons. The spike width was measured as the trough-to-peak width of the mean wide-band spike waveform. A representative waveform is shown. Putative MSNs were defined as units with > 0.6-ms spike width (green shaded areas). (C) Proportions of neurons classified as MSNs, fast spiking interneurons (28 and 25 neurons for the aDLS and pVLS, respectively), tonically active neurons (1 and 2 neuron (s) for the aDLS and pVLS, respectively), and unclassified interneurons (14 and 9 neurons for the aDLS and pVLS, respectively). Data are indicated as the mean ± s.e.m., and individual data are overlaid. Abbreviations: FSIs, fast spiking interneurons; TANs, tonically active neurons; and UIs, unclassified interneurons.

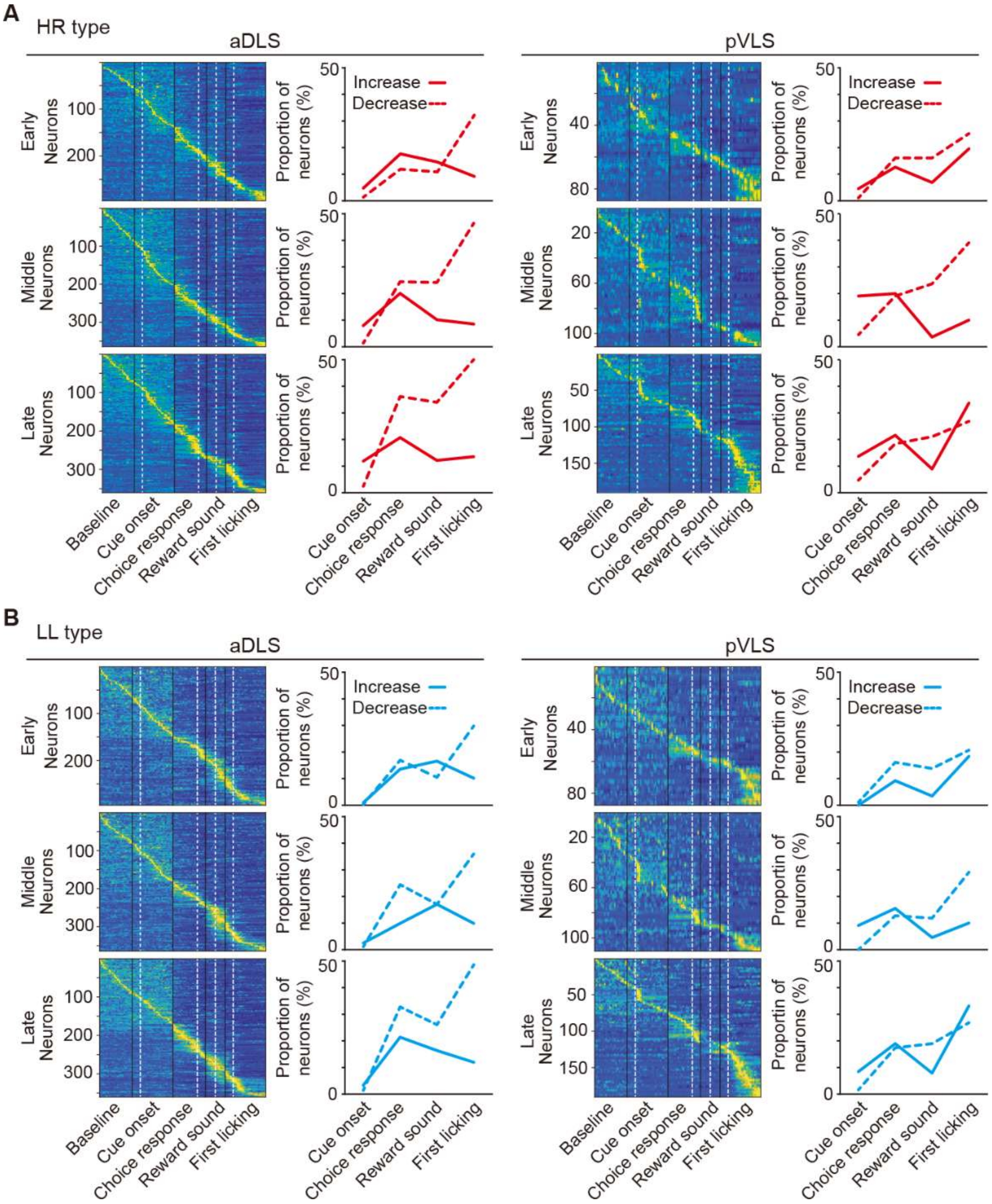


**Figure 5–figure supplement 3. Changes in event-related firing activity of HR and LL type neurons during the acquisition phase of auditory discrimination task.** (A and B) Pseudocolor maps indicate the z-scored event-related firing rate of HR (A) and LL (B) type neurons in the aDLS and pVLS. Spike data for each 50-ms bin during the baseline and time windows around the cue onset, choice response, reward sound, and first licking were concatenated in series. The proportion of neurons with increased or decreased activity is shown on the right side of the corresponding maps.

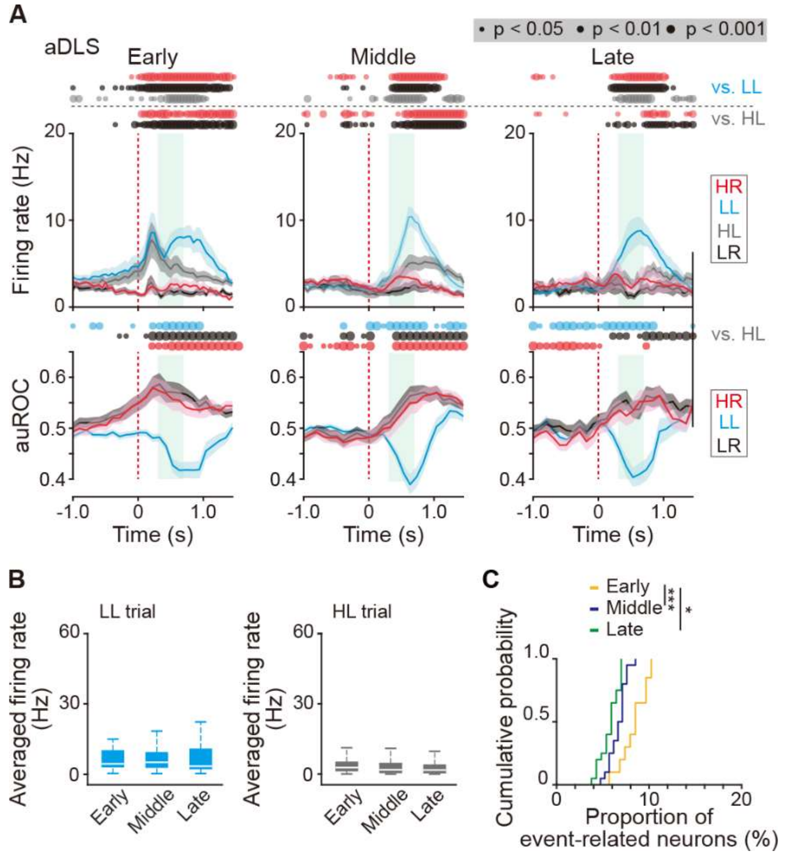


**Figure 5–figure supplement 4. Firing activity of reward sound-related LL type neurons in the aDLS.** (A) Mean firing rate (top rows) and auROC values (bottom rows) of reward sound-related LL type neurons in the aDLS at the early ($n$ = 49 neurons), middle ($n$ = 62 neurons), and late ($n$ = 59 neurons) stages. These data are aligned to the choice response. Green shadows show the period when the reward sound is presented (500 ± 200 ms after the lever press). Time bins with significant differences between the mean firing rate in the LL or HL trial and either of the rate in other three trials (Wilcoxon signed rank test) or the distribution of auROC values and 0.5 (Wilcoxon rank-sum test) are represented by the circles at the top. (B) Averaged firing rate during the reward sound period in the LL and HL trials at the early, middle, and late stages (Kruskal-Wallis test, LL trial, $\chi^2$ = 2.467, $p$ = 0.291, HL trial, $\chi^2$ = 4.426, $p$ = 0.109). (C) Cumulative probability of the proportion of the reward sound-related LL type neurons at the three stages (Kruskal-Wallis test, $\chi^2$ = 26.419, $p$ = $1.8 \times 10^{-6}$, *post hoc* Tukey–Kramer test, $p = 8.9 \times 10^{-7}$ for early vs. middle; $p$ = 0.010 for early vs. late; and $p$ = 0.067 for middle vs. late). Data are indicated as the mean ± s.e.m. (A) or the median and quartiles with the maximal and minimal values (B). *$p < 0.05$ and ***$p < 0.001$.

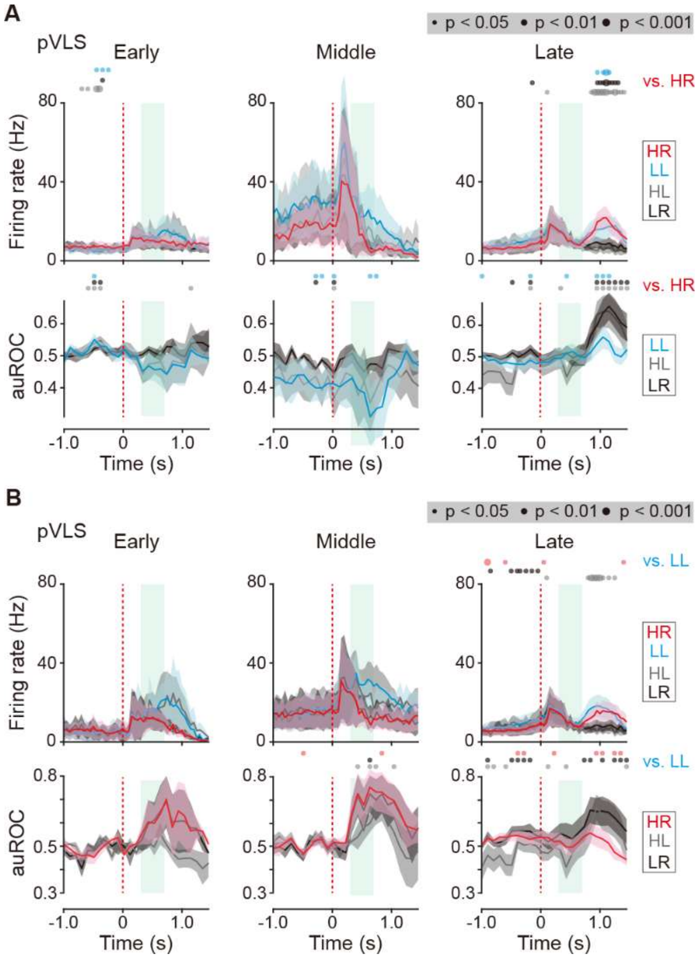


**Figure 5–figure supplement 5. Firing activity of. reward sound-related HR and LL type neurons in the pVLS.** (A) Mean firing rate (top rows) and auROC values (bottom rows) of the reward sound-related HR type neurons in the pVLS at the early (*n* = 6 neurons), middle (*n* = 4 neurons), and late (*n* = 17 neurons) stages. (B) Mean firing rate (top rows) and auROC values (bottom rows) of the reward sound-related LL type neurons in the pVLS at the early (*n* = 3 neurons), middle (*n* = 5 neurons), and late (*n* = 15 neurons) stages. These data are aligned to the choice response. Green shadows show the period when the reward sound is presented. Time bins with significant differences between the mean firing rate in the HR (A) or LL (B) trial and any of the rate in other three trials (Wilcoxon signed rank test) or the distribution of auROC values and 0.5 (Wilcoxon rank-sum test) are represented by the circles at the top. Data are indicated as the mean ± s.e.m.

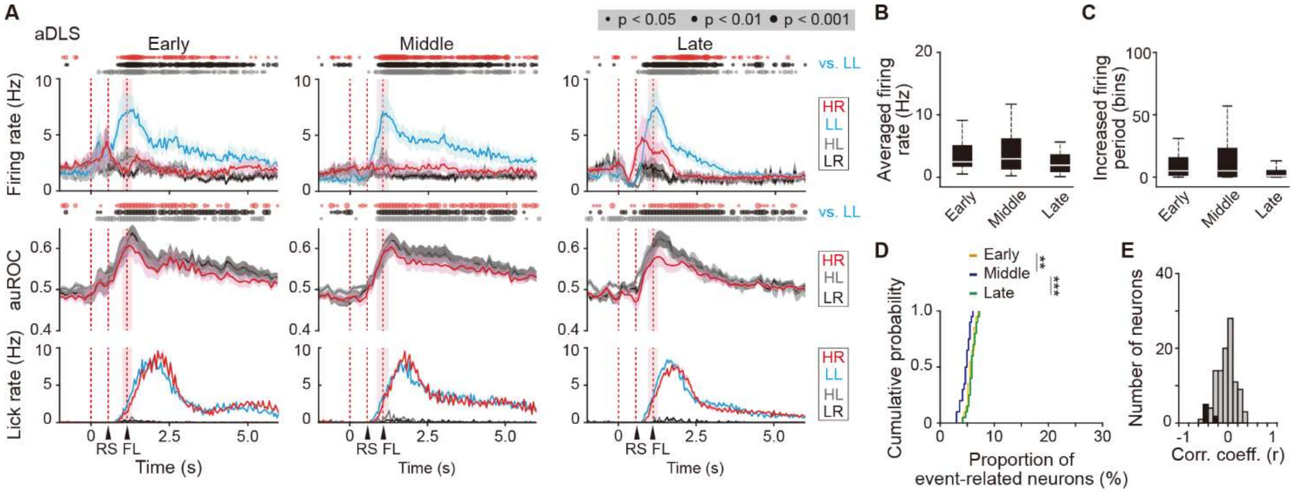


**Figure 6–figure supplement 1. Sustained activity after the reward of first licking-related LL type neurons in the aDLS**. (A) Firing rate (top rows), auROC (middle rows) and licking rate (bottom rows) after the reward of first licking-related LL type neurons in the aDLS at the early (*n* = 30 neurons), middle (*n* = 36 neurons), and late (*n* = 43 neurons) stages. These data are aligned to the choice response. Timings of the reward sound and first licking are shown. Time bins with significant differences between the mean firing rate in the LL trial and any of the rate in other three trials (Wilcoxon signed rank test) or the distribution of auROC values and 0.5 (Wilcoxon rank-sum test) are represented by the circles at the top. (B) Averaged firing rate for 5 s after the first licking in the LL trial (Kruskal-Wallis test, $\chi^2 = 4.474$, $p = 0.107$). (C) Averaged total number of time bins above 3 s.d. of the baseline firing for 5 s after the first licking in the LL trial (Kruskal-Wallis test, $\chi^2 = 6.306$, $p = 0.043$, *post hoc* Tukey-Kramer test, early vs. middle, $p = 0.989$, early vs. late, $p = 0.081$, middle vs. late, $p = 0.089$). (D) Cumulative probability of the proportion of first licking-related LL type neuron number (Kruskal-Wallis test, $\chi^2 = 18.952$, $p = 7.7 \times 10^{-5}$, *post hoc* Tukey-Kramer test, early vs. middle, $p = 0.003$, early vs. late, $p = 0.671$, middle vs. late, $p = 1.1 \times 10^{-4}$). (E) Distribution of the correlation coefficient between the numbers of spikes and licking. Closed column indicates the number of neurons showing significant differences between the two parameters (7 out of 109 neurons). Data are indicated as the mean ± s.e.m. (A) or the median and quartiles with the maximal and minimal values (B, C). $^{**}p < 0.01$ and $^{***}p < 0.001$. Abbreviations; RS, reward sound; and FL, first licking.

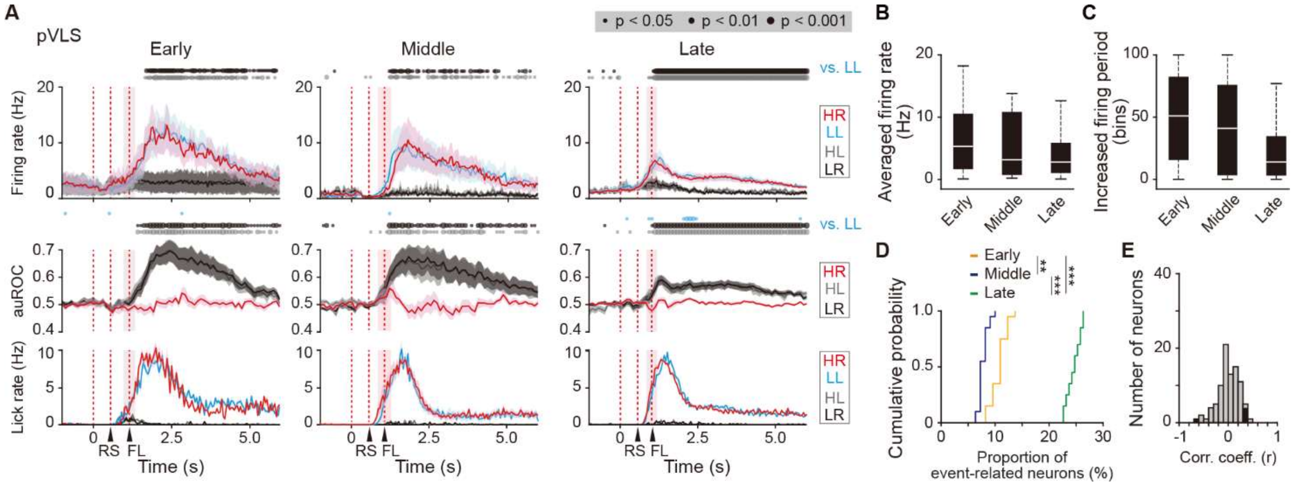


**Figure 6–figure supplement 2. Sustained activity after the reward of first licking-related LL type neurons in the pVLS**. (A) Firing rate (top rows), auROC (middle rows) and licking rate (bottom rows) after the reward outcome of first licking-related LL neurons in the pVLS at the early ($n$ = 16 neurons), middle ($n$ = 11 neurons), and late ($n$ = 63 neurons) stages. These data are aligned to the choice response. Time bins with significant correlations between the rate in the LL trial and either of the rate in other trials (Wilcoxon signed rank test) or the distribution of auROC values and 0.5 (Wilcoxon rank-sum test) are represented by the circles at the top. The timings of the reward sound and first licking are shown. (B) Averaged firing rate of pVLS neurons after the first licking in the LL trial (Kruskal-Wallis test, $\chi^2 = 2.327$, $p = 0.312$). (C) Averaged total number of time bins above 3 S.D. of the baseline firing of pVLS neurons after the first licking in the LL trial (Kruskal-Wallis test, $\chi^2 = 4.736$, $p = 0.094$). (D) Cumulative probability of the proportion of first licking-related LL type neuron number (Kruskal-Wallis test, $\chi^2 = 51.215$, $p = 7.6 \times 10^{-12}$, *post hoc* Tukey-Kramer test, early vs. middle, $p = 0.002$, early vs. late, $p = 5.1 \times 10^{-4}$, middle vs. late, $p = 9.6 \times 10^{-10}$). (E) Distribution of the correlation coefficient between the number of spikes and the licking. Closed column indicates the number of neurons showing significant correlations between the two parameters (5 out of 90 neurons). Data are indicated as the mean ± s.e.m. (A) or the median and quartiles with the maximal and minimal values (B, C). **$p < 0.01$ and ***$p < 0.001$. Abbreviations; RS, reward sound; and FL, first licking.

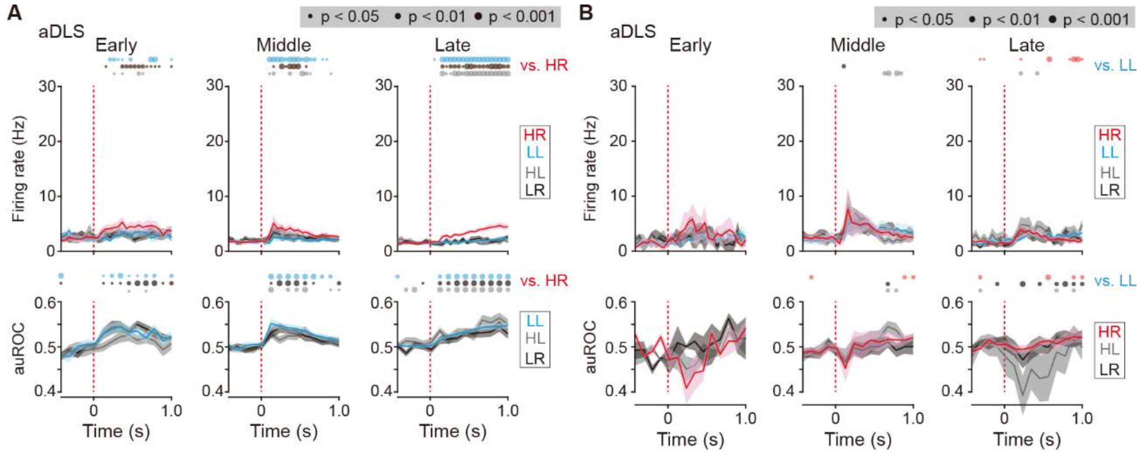


**Figure 7–figure supplement 1. Firing activity of cue onset-related HR and LL type neurons in the aDLS.** (A) Mean firing rate (top rows) and auROC values (bottom rows) of cue onset-related HR type neurons in the aDLS at the early (*n* = 14 neurons), middle (n = 29 neurons), and late (*n* = 43 neurons) stages. (B) Mean firing rate (top rows) and auROC values (bottom rows) of aDLS cue onset-related LL type neurons at the early (*n* = 3 neurons), middle (*n* = 9 neurons), and late (*n* = 12 neurons) stages. These data are aligned to the cue onset. Time bins with significant differences between the mean firing rate in the HR (A) or LL (B) trial and any of the rate in other three trials (Wilcoxon signed rank test) or the distribution of auROC values and 0.5 (Wilcoxon rank-sum test) are represented by the circles at the top. Data are indicated as the mean ± s.e.m. Abbreviation: RS, reward sound.

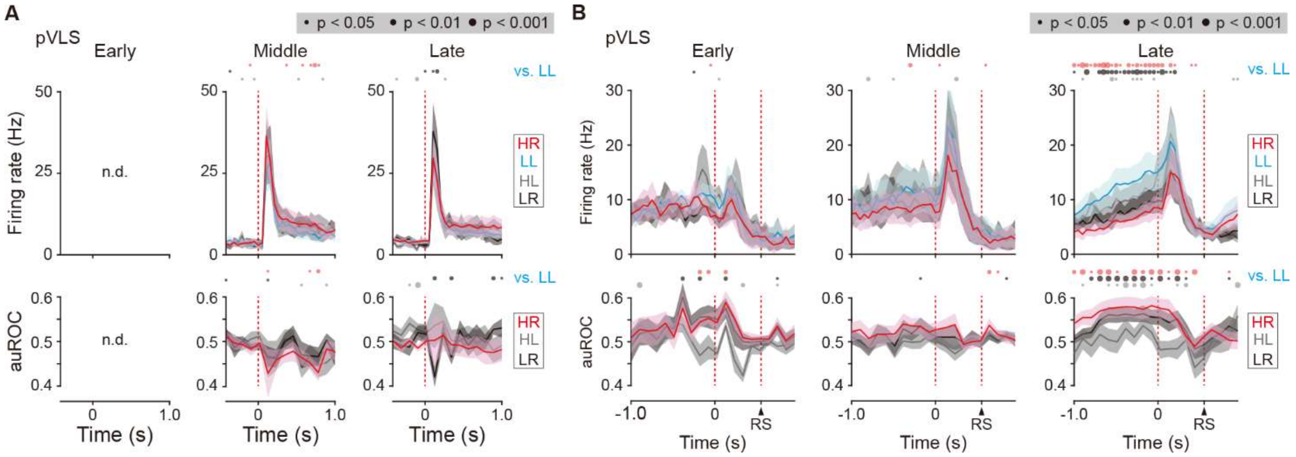


**Figure 7–figure supplement 2. Firing activity of cue onset- and choice response-related LL type neurons in the pVLS**. (A) Mean firing rate (top rows) and auROC values (bottom rows) of pVLS cue onset-related LL type neurons at the middle ($n$ = 10 neurons) and late ($n$ = 16 neurons) stages. (B) Mean firing rate (top rows) and auROC values (bottom rows) of pVLS choice response-related LL type neurons at the early ($n$ = 8 neurons), middle ($n$ = 17 neurons), and late ($n$ = 36 neurons) stages. The data are aligned to the cue onset (A) or choice response (B). The timing of the reward sound is indicated (B). Time bins with significant differences between the mean firing rate in the LL trial and any of the rate at other three trials (Wilcoxon signed rank test) or the distribution of auROC values and 0.5 (Wilcoxon rank-sum test) are represented by the circles at the top. n.d., not detected. Data are indicated as the mean ± s.e.m. Abbreviation: RS, reward sound.

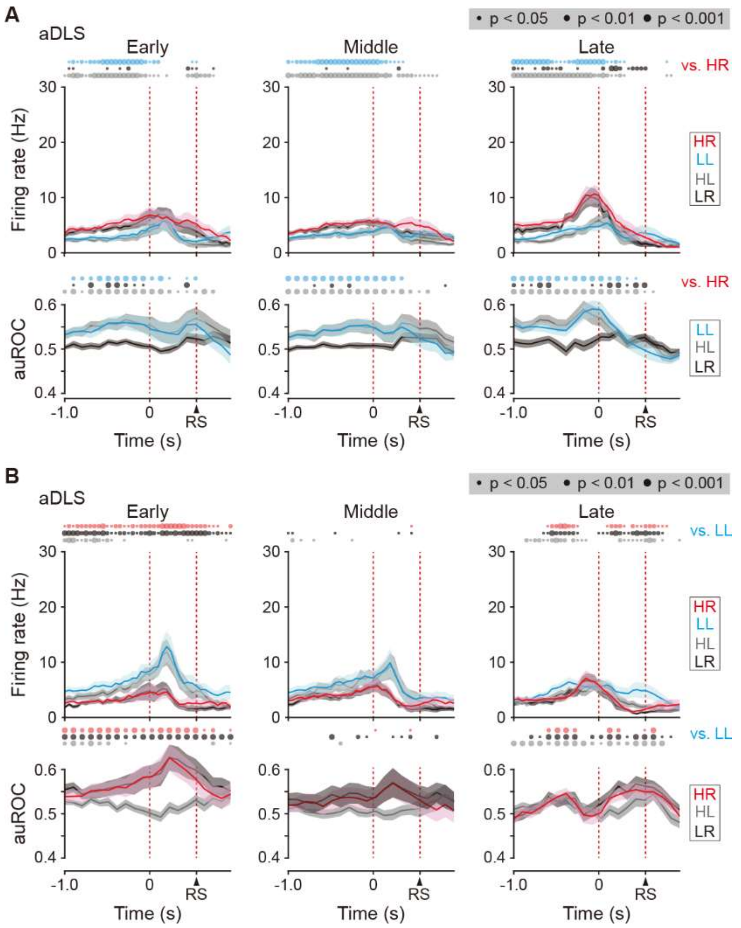


**Figure 7–figure supplement 3. Firing activity of choice response-related HR and LL type neurons in the aDLS.** (A) Firing rate (top rows) and auROC values (bottom rows) in aDLS choice response-related HR type neurons at the early (*n* = 52 neurons), middle (*n* = 73 neurons), and late (n = 75 neurons) stages. (B) Firing rate (top rows) and auROC values (bottom rows) in the aDLS choice response-related LL type neurons at the early (*n* = 40 neurons), middle (*n* = 36 neurons), and late (*n* = 77 neurons) stages. These data are aligned to the choice response. Time bins with significant differences between the mean firing rate in the HR or LL trial and any of the rate in other three trials (Wilcoxon signed rank test) or the distribution of auROC values and 0.5 (Wilcoxon rank-sum test) are represented by the circles at the top. Data are indicated as the mean ± s.e.m. Abbreviation: RS, reward sound.
